# Macroevolution of fly wings proceeds along developmental lines of least resistance

**DOI:** 10.1101/2025.01.09.632237

**Authors:** Patrick T. Rohner, David Berger

## Abstract

Evolutionary change necessitates genetic variation, and a reigning paradigm in biology is that rates of microevolution can be predicted from estimates of available genetic variation within populations. However, the accuracy of such predictions should decay on longer evolutionary timescales, as the influence of genetic constraints diminishes. Here, we show that intrinsic developmental variability and standing genetic variation in wing shape in two distantly related flies, *Drosophila melanogaster* and *Sepsis punctum*, are aligned and predict deep divergence in the dipteran phylogeny, spanning >900 taxa and 185 My. This alignment cannot easily be explained by constraint hypotheses unless most of the quantified standing genetic variation is associated with deleterious side-effects and effectively unusable for evolution. However, phenotyping of 71 genetic lines of *S. punctum* revealed no covariation between wing shape and fitness, lending no support to this hypothesis. We also find little evidence for genetic constraints on the pace of wing shape evolution along the dipteran phylogeny. Instead, correlational selection related to allometric scaling, simultaneously shaping developmental bias and deep divergence in fly wings, emerges as a potential explanation for the observed alignment. This suggests that pervasive natural selection has the potential to shape developmental architectures of some morphological characters such that their intrinsic variability predicts their long-term evolution.

## Introduction

A central aim in evolutionary biology is to predict evolution. Quantitative genetic approaches have played a central role in this endeavor by leveraging within-population estimates of evolvability in form of standing genetic variation and de novo mutational variation in quantitative traits to predict their evolution [1–4]). The translation of mutational variation at the nucleotide level into variation at the level of the phenotype is governed by developmental processes. Frequently, these processes channel random nucleotide changes into non-random phenotypic variation, giving rise to developmental bias [5–7]. These biases are recognized to in turn impact future adaptation by generating abundant substrate for evolution along certain phenotypic dimensions, while limiting it in others [5, 6, 8–11]. However, much controversy surrounds the timescale on which these biases constrain evolution. In particular, while limited genetic variation is predicted to slow down evolution, it is not expected to prevent phenotypic change entirely and estimates of evolvability within single populations are therefore expected to be poor predictors of macroevolutionary diversification [12–14].

Yet, several recent studies have challenged this standard expectation by showing correlations between evolvability estimates within populations and rates of macroevolution [15–18]. One line of evidence comes from studies on morphological evolution where mutational and standing genetic variation in morphological traits predict their long-term divergence [16, 18–20]. Due to the timescales over which evolution was observed, these relationships are hard to reconcile with the sole action of genetic constraints. Alternatively, it has been suggested that such correlations could result from natural selection that shapes the phenotypic effects of de novo mutations [4, 6, 11, 21–24]. According to this hypothesis, stabilizing selection moulds development so that deleterious effects of segregating genetic variants become reduced [24, 25] while the phenotypic effects of alleles under persistent directional or fluctuating selection instead become magnified [26–28]. In this process correlational selection acts on specific trait combinations so that developmental bias evolves, with the result that fitness-reducing phenotypic outcomes of mutations may become less frequent than expected by random chance and large mutational effects may mirror previous macroevolutionary divergence.

If past forces of selection indeed bias the phenotypic effects of de novo mutations, this would imply that the causal relationships between the processes of mutation, selection, and adaptation are more intricate than often assumed under standard models of evolution, with important implications for our ability to predict future evolution from current quantitative genetic parameters [6, 22, 28, 29]. However, the role of selection in shaping mutational effects (i.e., development biases) remains controversial and has been disputed on theoretical grounds [25, 26, 30–36], and reconciling the observed relationships between evolvability and macroevolution with processes occurring at microevolutionary scales remains a fundamental challenge [19, 20, 29, 37–39].

Here we address this controversy by extending recent analyses on the relationship between developmental bias and evolutionary divergence in dipteran wings. Houle et al. [18] demonstrated that mutations cause nonrandom phenotypic variation in the wings of *Drosophila melanogaster* and, astonishingly, predict 40 My of divergence across the Drosophilidae. These results were recently complemented by a study [16] showing that developmental bias in sepsid flies, a clade that diverged from the Drosophilidae around 60 Mya, is related to the mutation bias and macroevolutionary patterns observed by Houle et al [18]. Here we show that this alignment holds on even larger timescales, providing evidence for a relationship between *de novo* mutational input and macroevolution that unfolded over 185 My. We show that the genetic constraint hypothesis alone is a poor fit to the observed patterns. Alternatively, correlational selection on wing traits as a causative agent shaping both developmental bias and deep divergence remains a plausible, yet disputed, explanation for the observed patterns. Irrespective of their ultimate explanation(s), our findings show that deep divergence in dipteran wings can be reasonably well predicted from their intrinsic developmental variability, even when such variability may fail to predict evolution on shorter timescales. This challenges our understanding of the fundamental processes that govern the emergence and evolution of phenotypic variation.

## Results

Developmental bias can be quantified by studying how genetic or environmental perturbations affect developmental outputs in the form of phenotypic variation. One way of doing so is to study how phenotypic variation in multivariate characters is generated by *de novo* mutation, captured by the mutational variance-covariance matrix, **M**, an approach utilized by Houle et al. [18] to capture mutational bias in wing shape in *D. melanogaster*. An alternative way of quantifying developmental variability is to measure fluctuating asymmetry between left and right homologs of paired bilateral structures. Because the left and right sides of the same organism share the same genome and environment, differences between bilateral homologues can be attributed to developmental noise, and differences in the degree of fluctuating asymmetry among phenotypes thus serve as a measure of bias in the developmental program [40, 41]. This approach was used by Rohner & Berger [16] to capture the developmental covariance matrix, **D**, for wing shape in sepsid flies. This covariance matrix thus captures how traits (co)vary in response to random developmental perturbations. Here we first use these previous estimates of **M** (by [18]) and **D** (by [16]) to predict 185 My of macroevolution across 43 families and >900 species of Diptera. We then evaluate competing hypotheses invoking genetic constraints and correlational selection to reconcile the observed alignment between the generation of *de novo* variation and deep macroevolution.

We focus on the evolution of wing shape within the Eremoneura, a clade within the higher flies (Brachycera) that is about 185 My old [42] and contains more than 64,000 species (including, among others, the fruit, flesh, house, and tsetse flies). To quantify variation in landmark positioning (see Supplementary Data Fig. 1, Supplementary Data Table 1) within and between families, we took advantage of published scientific illustrations and pictures of fly wings from the taxonomic and systematic literature (e.g. [43, 44], see Fig. 1, Supplementary Data Table 2). Our final dataset contained 988 observations from 933 different species. To first assess the accuracy of using illustrations (n = 414) compared to pictures of wings (n = 574), we computed the partial least-squares (PLS) correlation between coordinates derived from illustrations and pictures for 19 species where both types of data were available. This correlation was very strong and statistically significant (r_PLS_ = 0.98, Z = 4.21, P < 0.001, see Fig. 1, Supplementary Data Fig. 2). Similarly strong associations were found when comparing the average wing shape for each genus based on illustrations with the average shape calculated based on measurements taken from pictures (r_PLS_ = 0.93, Z = 6.35, P < 0.001, n = 64), showing that shape information derived from illustrations adequately captures shape variation derived from photographs.

**Figure 1:**
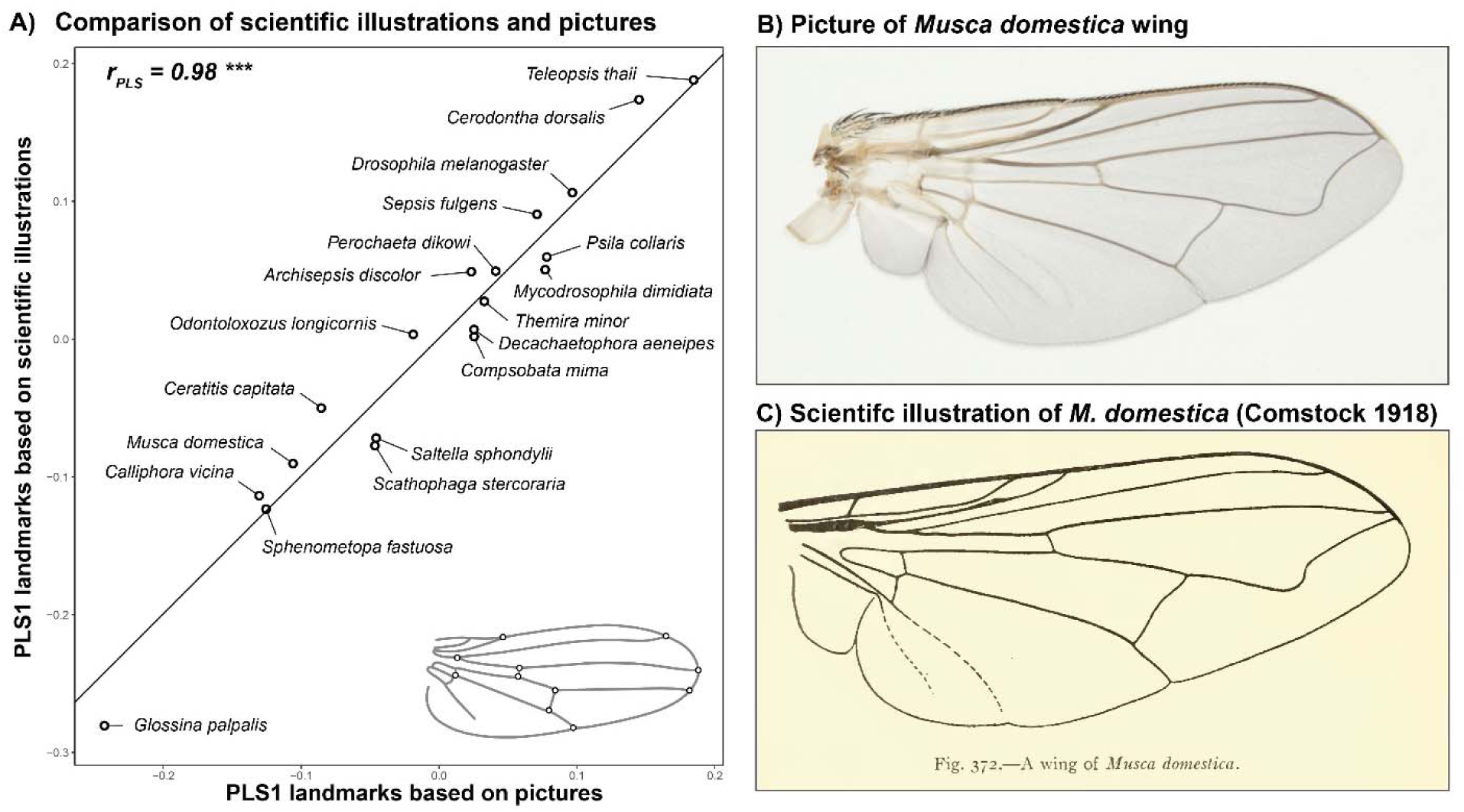
Partial least squares plot showing a strong correlation between the shape measurements derived from pictures or scientific illustrations for 19 species where both sources of data were available (A). Panels B) and C) highlight the similarity between a picture of the wing of a house fly (*Musca domestica*) and an illustration of a wing of the same species (the latter modified from [104]).

### Developmental and mutational variance in wing shape is correlated with its deep macroevolutionary divergence

The 43 fly families in our dataset that had five or more representatives differed strongly in their wing shape (Fig. 2; Procrustes ANOVA: F_42,926_ = 49.60, Z = 27.74, P <.001, R^2^ = 0.73). Leave-one-out cross-validation led to a correct classification of 82.3% of all individuals (Canonical Variate Analysis), indicating that fly families can be differentiated based on wing shape (see Supplementary Data Table 3, Supplementary Data Fig. 3). To quantify the macroevolutionary dynamics of wing shape, we computed the evolutionary rate matrix **R** based on the inverse of the phylogenetic relationship matrix among dipteran families [45] using animal models in *ASReml-R* [46]. Because large species-level phylogenies are lacking on this broad phylogenetic scale, we based our analysis on a recent phylogeny that leveraged transcriptomes (3,145 genes) to resolve the phylogenetic placement among families [47] (see Fig. 1). The species represented in our database that fall within these families were treated as replicated measures for each family’s wing shape at the tip of the phylogeny. Again, we only included the 43 families represented by at least 5 species in our analyses. Macroevolutionary divergence was mostly related to the relative positioning of the first branch of the radial vein (R_1_) and the placement of the two cross-veins along the proximo-distal axis (see Fig. 2).

**Figure 2:**
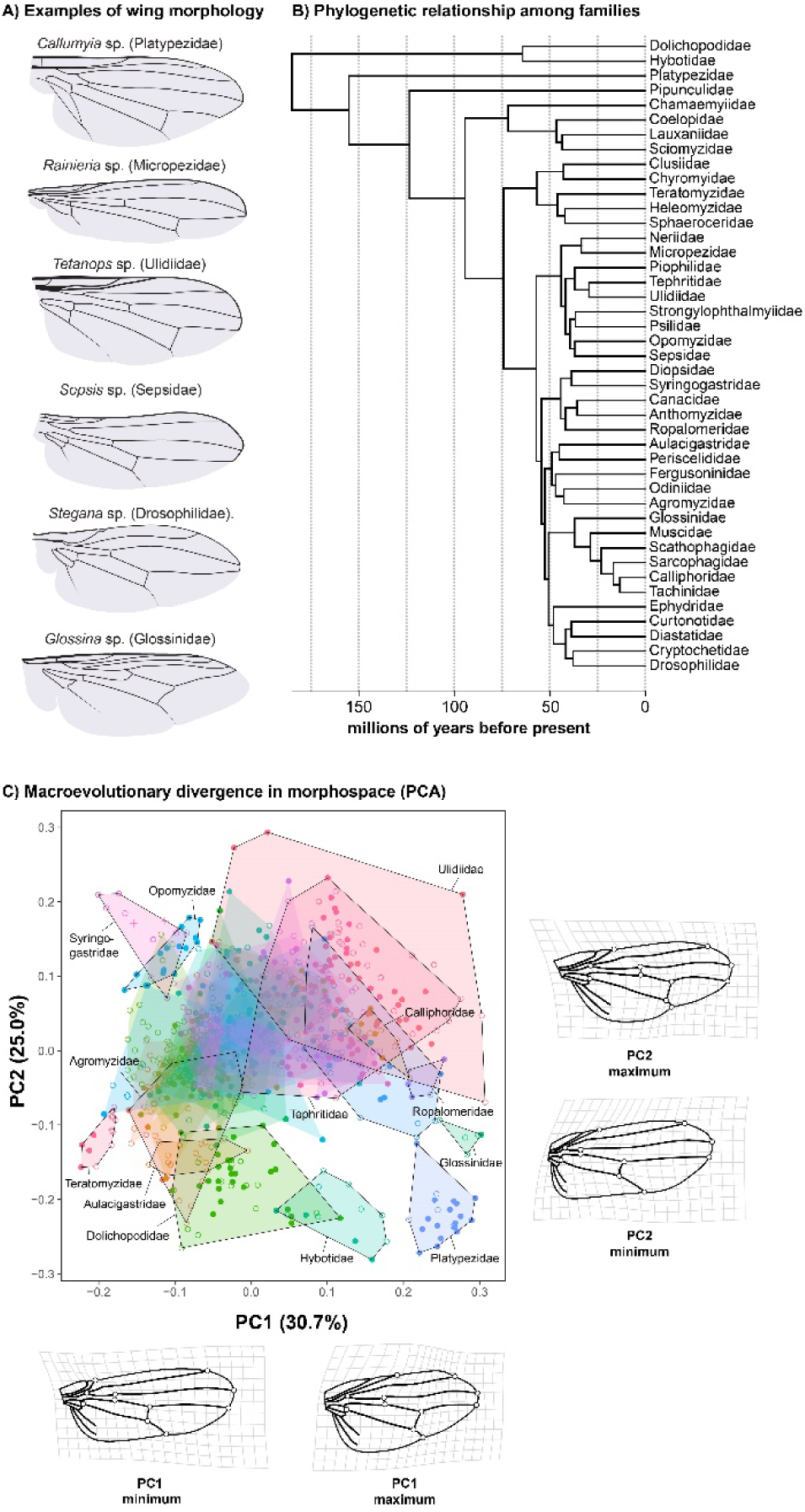
Although fly wings evolve slowly, there is large macroevolutionary divergence in wing shape (examples highlighted in panel A). Panel B shows the evolutionary relationships among the different families within the Eremoneura [47]. The phylogeny was calibrated using the approximate age of the Eremoneura [42], as well as the split between Drosophilidae, Muscidae, and Tephritidae [94], respectively. Panel C shows an evolutionary morphospace defined by the first two principal components. Individuals are grouped by family (hulls). An arbitrary set of families is highlighted. Families with less than five observations were excluded from the plot. Shape deformations associated with minimal and maximal loadings relative to the average wing shape indicated with deformation grids for the first two PCs. To improve visibility, the magnitude of shape changes was reduced by a factor of 0.5. The outline of the wing was based on a drawing of *Camptoprosopella vulgaris* (Lauxaniidae) after Curran 1934 [105].

Next, we tested whether deep divergence among families is related to mutational and developmental bias observed in drosophilids and sepsids. Specifically, we compared **R** to the previously estimated **D** and **M** matrices in *S. punctum* and *D. melanogaster* using a modified version of Krzanowski’s common subspace analysis following the method described in [48] [also see 18, 49]. In brief, we compared the logarithmized variances of both matrices along the same set of orthogonal phenotypic dimensions of the wing. To limit bias in our estimates of effect sizes [48, 49], we chose to represent these phenotypic dimensions by the eigenvectors of an independently estimated third matrix —the phenotypic variance-covariance matrix, **P**— measured in *S. fulgens* (this is a morphologically distinct but relatively close relative of *S. punctum* placed in the same species group [50] within the same genus). For consistency, we use this matrix as the reference to generate comparisons of different variance-covariance matrices throughout this study. However, we also repeated all comparisons by using other matrices as reference (see Supplementary Data Table 4) which showed that our conclusions do not depend on the matrix chosen as reference.

To make sure that all matrices were compared along subspaces in which there was statistically verified variation, we estimated the rank of the matrices by employing factor analytical modelling using ASReml-R [46]. Matrices were then compared along the first *k* dimensions of **P**, with *k* equal to the rank of the matrix with the lowest rank (k = 10). We applied this approach for any pair of variance-covariance matrices compared in this study. If macroevolutionary divergence across dipteran families can be predicted by developmental bias, we expect **R** to show similar relative amounts of variation as **D** and **M** along the eigenvectors of **P**. Regressing the resulting (logarithmized) variances of **R** on the corresponding variances of **M** and **D**, we indeed find that the morphological variation representing macroevolutionary change is similar to that generated by mutation (**R** on **M**: slope = 0.66 [0.53, 0.77] 95%CI, r = 0.89 [0.78, 0.94], Fig. 3) and developmental perturbations (**R** on **D**: slope = 0.61 [0.5, 0.71]; r = 0.87 [0.78, 0.92], Fig. 3). If analysing all 18 phenotypic dimensions of the wing by also including the eight additional wing-dimensions for which we could not statistically certify significant variation at all biological levels compared, the relationships become even stronger (Supplementary Data Fig. 4). This suggests that macroevolutionary divergence among 43 dipteran families that unfolded over 185 My is aligned with developmental lines of least resistance and can thus — at least to some degree — be predicted from intrinsic developmental variability documented in single species. (For comparison, the age of the clade we study here predates the split between marsupials and placentals in the Jurassic [51]).

**Figure 3:**
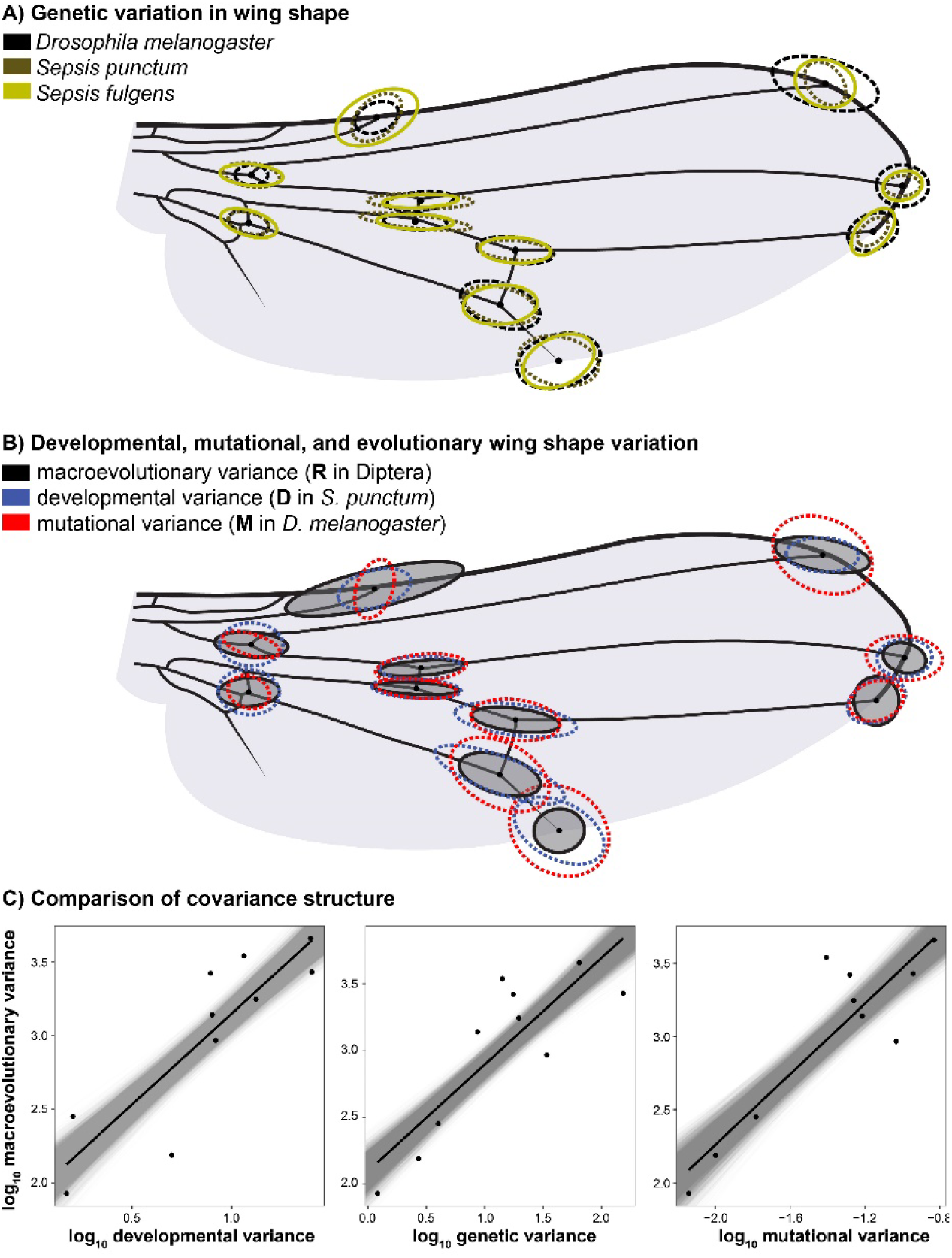
A) Similarity in standing genetic variation in wing node positioning across *Drosophila* and two sepsid flies that diverged ca. 64 Mya. The average location of each landmark was shifted to match a typical sepsid wing. B) Variation in wing shape due to developmental (**D**), mutational (**M**), and macroevolutionary (**R**) variation. Matrices shown in panels A and B were scaled by their trace to facilitate comparison. Panels in C) show the results of a common subspace analysis where the amount of developmental, standing genetic, and mutational variance predicts the macroevolutionary variance along the same set of orthogonal vectors (i.e., the first 10 eigenvectors of the phenotypic variance-covariance matrix estimated in *S. fulgens*). Grey lines indicate the distribution of regression slopes using REML-MVN resampling (n = 10,000).

### No evidence for deleterious pleiotropy constraining the evolution of wing shape

The relationship between developmental bias and macroevolution is consistent with fundamental constraints of wing shape development and evolution. However, due to their polygenic basis and large mutational target sizes, the evolution of quantitative characters is typically not expected to be strongly constrained over the long timeframes studied here [2, 20, 52]. Indeed, the study on drosophilids by Houle et al. [18] found that a lack of mutational input is unlikely to explain the alignment between **M** and **R** in drosophilid wing evolution. To explore the possible influence of genetic constraints on the studied macroevolution of wing shape, we calculated the expected amount of divergence along the 10 analyzed wing shape dimensions (see Fig. 3) under a scenario of pure genetic drift, which predicts that the rate of divergence should correspond to two times the mutational variance per generation [53]. Thus, if genetic constraints are limiting the evolution of some wing dimensions, we expect that the observed rates of divergence should be approximated by the predicted divergence based on the rate of mutational input. However, assuming an average of a single fly generation per year, and basing our calculations on estimates of **M** in *D. melanogaster* [18], we find that the observed macroevolutionary variance along each of the 10 dimensions is around 10^4^ times smaller than expected under drift (Supplementary Data Table 5). Because most species studied here undergo more than one generation per year, this calculation underestimates the expected divergence under drift (for instance, central European populations of *S. punctum* have at least four generations per year, and *D. melanogaster* has about 15 generations per year [54, 55]). Thus, genetic constraints alone are unlikely to explain the low rates of divergence and the observed correlation between developmental bias and macroevolution.

The constraint hypothesis would remain viable if most of the quantified mutational variation has deleterious pleiotropic side-effects on other unmeasured traits, rendering the variation effectively unusable for adaptive evolution [18]. We tested this hypothesis by quantifying genetic variation in fitness-related traits and wing shape in *S. punctum*. If wing shape evolution is indeed constrained by deleterious pleiotropy, we expect to find genetic covariation between wing shape and fitness components that are functionally unrelated to wing shape or flight. Rearing the 71 iso-female lines of *S. punctum* assayed for wing shape [16] in a common garden experiment limiting direct selection on flight (flies were kept in 50ml glass vials), we found significant heritable variation among iso-female lines in adult longevity (*Χ*^2^_(1)_ = 8.62, P = 0.003), developmental rate (*Χ*^2^_(1)_ = 369.62, P <.001), juvenile survival (*Χ*^2^_(1)_ = 89.79, P <.001), as well as body size (*Χ*^2^_(1)_ = 225.76, P <.001) but not in early reproductive success (*Χ*^2^_(1)_ = 0.61, P = 0.218). About half of all pairwise genetic correlations between fitness components based on best linear unbiased predictors (BLUPs) were positive and statistically significant, indicating that some lines had an overall higher fitness than others. For instance, iso-female lines with high fecundity also had a faster developmental rate (t_(70)_ = 3.74, r = 0.41 [0.20, 0.59] 95%CI, P <.001), larger adult size (t_(70)_ = 2.97, r = 0.34 [0.11, 0.53], P = 0.004), and longer adult lifespan (t_(70)_ = 2.44, r = 0.28 [0.05, 0.48], P = 0.017; Supplementary Data Fig. 5). This collinearity was also reflected by all five fitness components loading in the same direction on the dominant principal component (PC1) describing trait variation (Fig. 4A). Because individuals with high scores on PC1 had higher fitness across all fitness components and considering that PC1 also explained a larger proportion of the total variation than expected by chance (37.1%; P_RAND_<.001, see Fig. 4B), these patterns suggest that PC1 captures deleterious pleiotropic alleles affecting life-history traits and variation in overall genetic quality. If wing shape is associated with deleterious side-effects, one would thus expect wing shape to covary with PC1. However, PC1 was not related to genetic variation in wing shape (r_PLS_ = 0.28, Z = 0.14, P = 0.445) and we also found no evidence for a relationship between any of the 5 fitness components and wing shape when all variables were simultaneously analyzed in a two-block partial least squares analysis (Fig. 3C, r_PLS_ = 0.39, Z = 0.54, P = 0.299). To test for stabilizing selection, we also estimated the correlation between the iso-female lines’ fitness and their multivariate residual from the mean wing shape. None of the correlations between wing shape residuals and the five fitness correlates were significant (see Fig. 4D; |r| < 0.21, P > .085). All these analyses were repeated while excluding a single outlier (see Supplementary Data Fig. 5) with the same result. Thus, based on these data, we find no support for the hypothesis that deleterious pleiotropy acts as an evolutionary constraint on wing shape divergence. We note, however, that our experiment would have had limited power to detect more subtle covariation with fitness that might still affect the evolutionary potential of wings.

**Figure 4:**
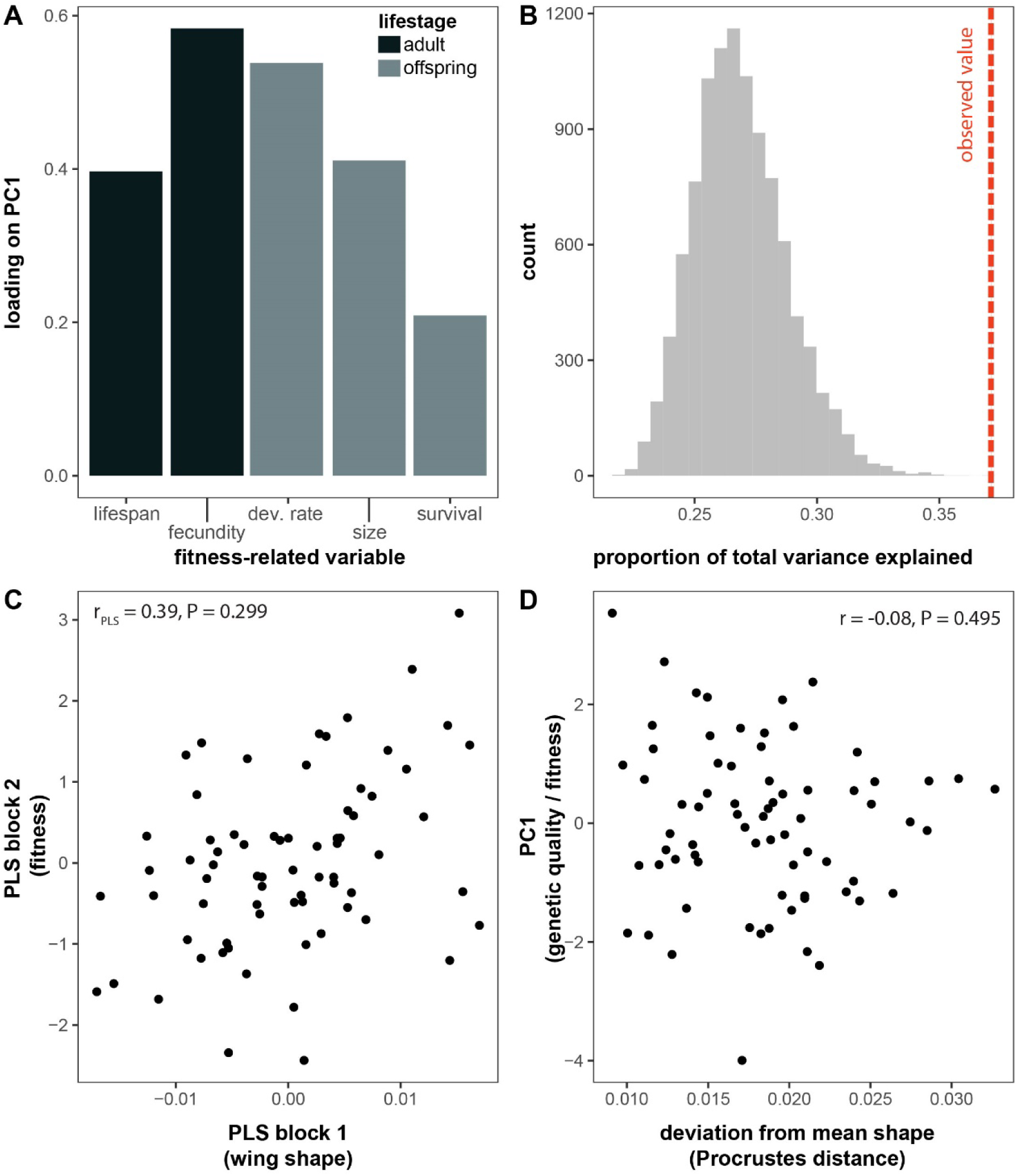
Loadings of fitness components measured in adults and offspring on PC1 all have the same sign (A), suggesting that different fitness components are correlated, indicating a main axis of genetic quality. PC1 explained more variation than expected by chance (B) (observed variance: red hatched line, null distribution: grey bars). However, despite significant genetic variation in wing shape and fitness, we found no relationship between wing shape and the five fitness components investigated (C). There was also no relationship between the isofemale lines’ fitness (here indicated as scores on PC1) and their multivariate distance to the average wing shape, as expected if wing shape was under stabilizing selection.

### Evolution is not faster along the wing dimensions with the greatest developmental variance

If developmental bias indeed acts as a constraint on evolution, we would also predict that macroevolutionary divergence along phenotypic dimensions with little developmental variance **D** to be relatively slow [1, 2, 4]. To test this prediction, we first reconstructed the evolutionary history of wing shape and quantified the alignment between developmental (**D**) or mutational (**M**) bias and the direction of evolutionary shape change on each branch of the phylogeny. Here, we quantified these alignments as the proportion of the trace of **D** (or **M**) that was captured by the shape change vector along individual branches. We then tested whether evolutionary shape changes more closely aligned with the main axes of **D** (or **M**) have been faster than wing shape changes unrelated to developmental bias. Contrary to the expectation under the constraint hypothesis, we found no strong correlation between evolutionary rate and the alignment of divergence with **D** (r = −0.23) or **M** (r = −0.15) (see Fig. 5B and Supplementary Data Fig. 6). The observed correlations are significantly lower than that expected under simulated Brownian Motion (both P_RAND_ < 0.01; Fig. 5C).

**Figure 5:**
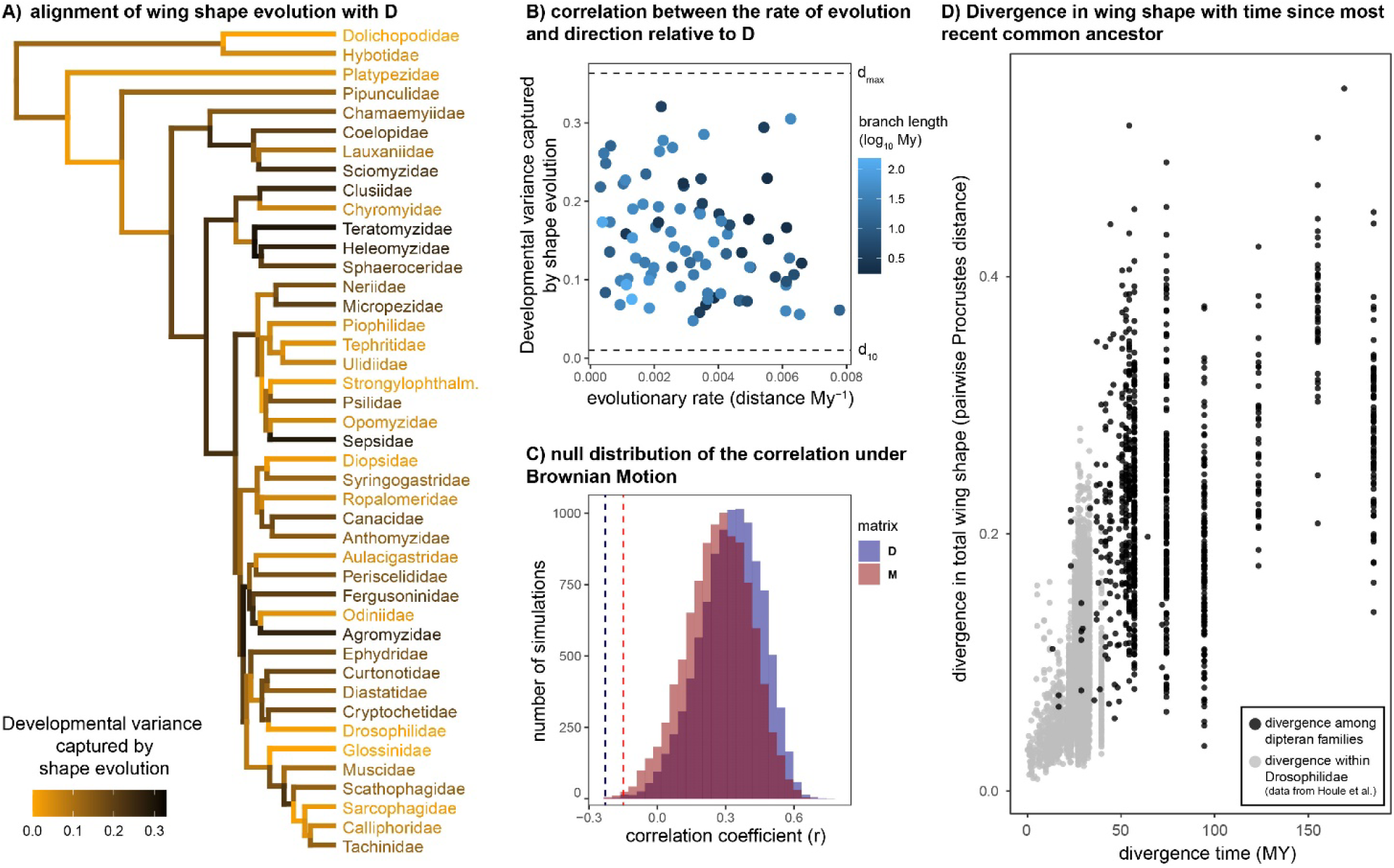
If developmental bias constrains evolution, morphological evolution is expected to be faster if it aligns well with the main axes of the **D** matrix. To test this hypothesis, we first quantified the vector of shape change along the edges of the phylogeny and computed the developmental variance captured by shape evolution. If evolution occurs mainly along the main axes of **D**, this variance will be large and close to the variance explained by the first eigenvector of **D** (d_max_), indicating an alignment between shape evolution and developmental bias (see panel A). We then correlated the strength of this alignment with the rate of evolution (shape evolution in Procrustes distance per My) along the branch, expecting a positive relationship if fast rates of evolution are constrained along wing dimensions with high variability. In contrast to this expectation, the rate of evolutionary change does not depend on its alignment with **D** (panel B) (or **M**, Supplementary Data Fig. 6). This observed correlation (hatched vertical lines in panel C) is also much lower than what is expected under pure Brownian Motion (panel C). Panel D shows the increase in wing shape divergence with phylogenetic distance. Data in grey (below 40 My) are from Houle et al [18].

Taken together, our results render a simple constraint hypothesis unsuited to explain the observed pattern of macroevolution, which is perhaps unsurprising given the implausibility of genetic drift or unidirectional selection on wing characters over such long timescales. Holstad et al. [19], recently reported a correlation between evolvability and macroevolutionary rates and argued that such patterns can emerge from rapid fluctuating selection around a global phenotypic adaptive zone defined by persistent stabilizing selection common to all taxa. In this scenario, evolving taxa are constantly tracking, but often lagging behind, rapid shifts in phenotypic optima. Such adaptive tracking would be more efficient for traits with abundant genetic variation, resulting in greater differentiation between taxa, whereas traits with low levels of variation would show greater lags and less differentiation, creating a positive correlation between trait evolvability and macroevolution while evolutionary divergence would remain overall low. However, Holstad et al. based their conclusions on patterns observed over a few million years (typical divergence times <1My), and as pointed out by the authors, the signal of fluctuating selection and associated tracking would likely get overridden by episodes of genetic drift and divergent selection operating over the long macroevolutionary timescales we study here ([19, 56], see also [57]). Indeed, our data on fly wings show the footprint of substantial accumulated divergence between lineages, tracing far back to deep splits in the dipteran phylogeny (Fig. 5D, Supplementary Data Fig. 7). Moreover, the fluctuating selection scenario does not by itself generate a strong phylogenetic signal in the data over time. In contrast, the phylogenetic signal in our data explained 86 percent of the macroevolutionary variance among families (mean phylogenetic heritability weighed by the total amount of species variance of each shape variable, see [18]). Thus, our results seem incompatible with the constraint hypothesis invoking fluctuating selection around stationary phenotypic optima envisioned by Holstad et al. [19].

### Correlational selection acting on multiple levels of biological organization?

An alternative explanation for the observed alignments emerges if we consider that different dimensions of the wing may experience different strengths of directional and stabilizing selection, such that developmental bias has itself evolved to align with the fitness surface [6, 24, 58, 59]. Under this scenario, proportionality between **D**, **M**, and **R** is observed, not because development constrains macroevolutionary rates, but because pervasive correlational stabilizing selection has restricted developmental variability, mutational effects, and divergence to occur along similar phenotypic dimensions. One such pervasive force is correlational selection for optimal allometric relationships between morphological characters [2, 39, 60]. Indeed, insect wings show strong allometric scaling, likely due to functional constraints [61, 62]. We therefore tested whether the observed patterns could be explained by allometric scaling so that the relationship between **R** and **D**/**M** could solely be ascribed to allometric scaling across Diptera.

Studying the subset of illustrations and pictures that had an associated scale bar (n = 127 species), we find evidence for interspecific allometry (multivariate regression of shape against log centroid size: F_1,126_ = 18.7, P <.001, R^2^ = 0.13) which correlates with the intraspecific wing shape allometry vector previously documented in *S. punctum* (vector correlation: r = 0.5, P <.001). This is consistent with studies suggesting conserved allometric scaling across Diptera [61]. This interspecific allometric vector captured more variation in both **D** and **R** than expected by chance (P_RAND_ <.001), indicating that correlational selection for optimal allometric scaling may be causally involved in shaping developmental bias and macroevolutionary divergence in wing shape (Fig. 6A). Interestingly, however, when recalculating the **R** matrix based on residual wing shape after the effect of a common allometric slope was removed, we still recover a strong alignment between developmental bias and evolutionary divergence (**M**: b = 0.71 [0.56, 0.83], r = 0.82 [0.68, 0.90]; **D**: b = 0.55 [0.42, 0.66], r = 0.79 [0.69, 0.88]). Hence, while our analysis provides indirect support for a role of correlational selection on allometric scaling in driving the alignment between developmental bias and deep divergence, the alignment persists even when controlling for the effect of allometry.

**Figure 6:**
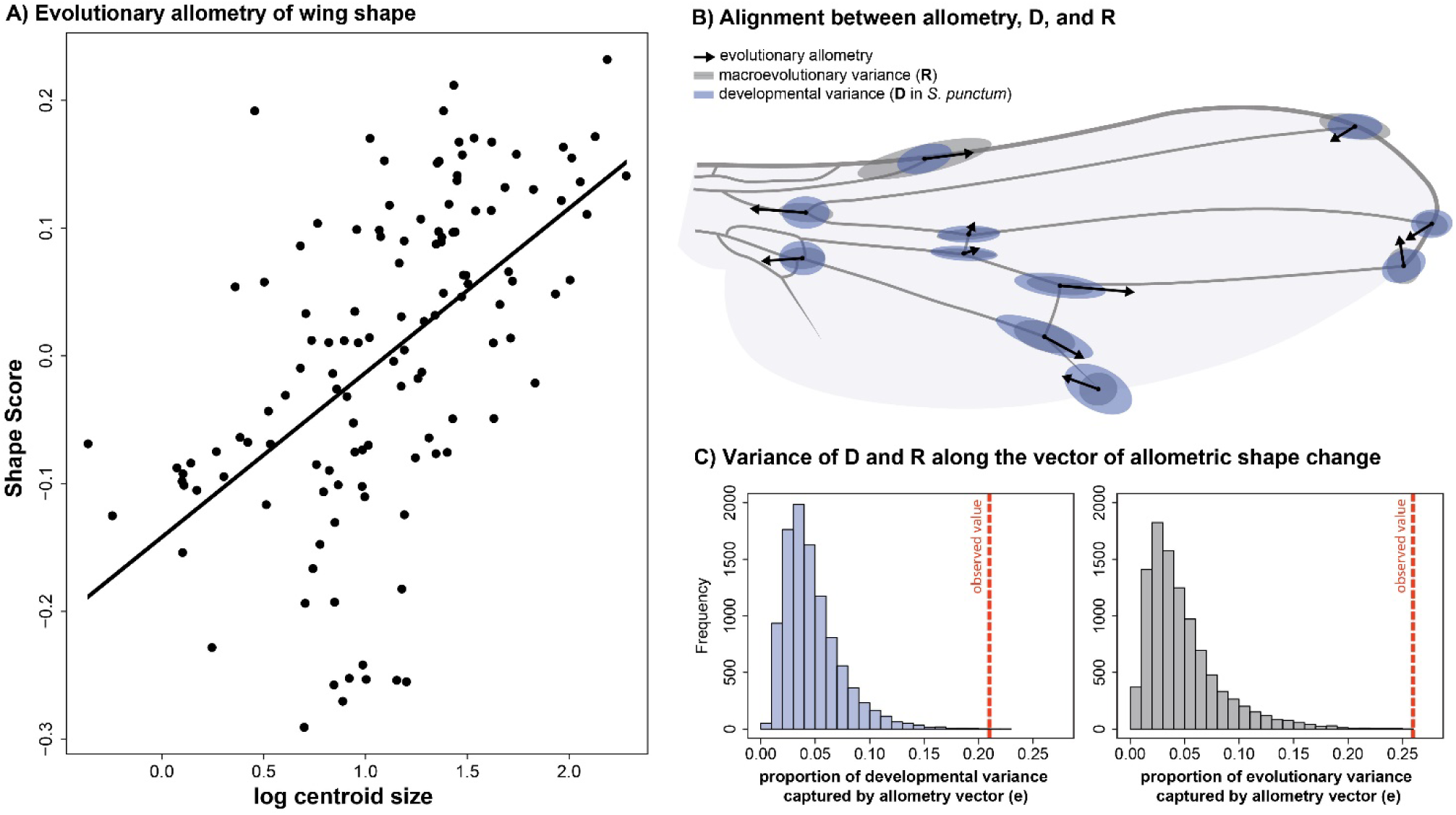
Wing shape changes with size across 127 species of flies (spanning 30 different families), indicating allometric scaling of shape (A). These evolutionary allometric shape changes align with the main axes of developmental and macroevolutionary variance in fly wings (B). These alignments are much stronger than expected by chance (C). The displacement of relative landmark positioning with an evolutionary increase in wing size is indicated by black arrows.

## Discussion

Here we leveraged within-species estimates of developmental (**D**) and mutational (**M**) variability to assess developmental bias and show that this bias can predict macroevolutionary diversification in deep time. Quantitative genetic theory rests firmly on the assumption that, due to genetic constraints on rates of evolution, accurate estimates of mutational and genetic covariance matrices can be used to predict evolutionary change in correlated morphological characters. However, much debate remains surrounding the utility of these approaches when applied on longer evolutionary time scales [4, 12, 14, 22, 24]

First, it remains uncertain if there is enough stability in the amount standing genetic variation in correlated characters (captured in the genetic variance-covariance matrix, **G** [2]) to allow accurate predictions of their long-term evolution [13]. Indeed, if selection and drift reshape **G**, then snapshots of standing genetic variation at any point in time are likely to be poor predictors of evolution, even over only a few hundred generations. In contrast to this notion, developmental bias (**D**, **M**, and **G**) in fly wings remains surprisingly conserved across the Drosophilidae and Sepsidae (Fig. 3A, B), clades that diverged from each other around 60 Mya. Similarly, McGlothlin et al. [63] showed that **G**, while having evolved across species of *Anolis* lizard, had retained its main dimensionality across >20 My of species divergence.

Second, however, even if **G** and **M** were to remain constant, predictability relies on the consistency of natural selection, which is likely to fluctuate even in the short term [64, 65]. It would thus seem that evolutionary prediction might be limited to special circumstances. Yet, alignments between standing genetic variation within populations and macroevolutionary rates over a couple of millions of years have been observed for morphological features in plants, insects, and vertebrates [4, 19, 20, 63, 66, 67], notably up to 40 My in *Anolis* lizards [63]. Here we show that the recently reported correlations for fly wings [16, 18] can extend even longer, with wing shape evolution being predictable —at least to some degree— over 185 My.

Macroevolution has been suggested to unfold at slow pace along genetic lines of least resistance delineated by the architecture of the developmental system [68]. While such patterns on their own are compatible with evolutionary constraints, they are hard to reconcile with observations of contemporary adaptation being exceedingly fast [69, 70], questioning the general applicability of genetic constraints as an explanation for evolutionary stasis. Indeed, several analyses have demonstrated that evolutionary stasis in the fossil record may not represent slow rates of evolution, but rather abundant adaptive change in response to fluctuating selection within certain boundaries [19, 37, 57, 71, 72]. This scenario is compatible with other observations of stasis in shape evolution in the fossil record despite episodes of strong directional selection [73] and was recently proposed to explain evolvability-macroevolution relationships on the scale of a couple of million years [19]. However, under this scenario, any macroevolutionary divergence due to drift or directional selection would be expected to erode such a relationship over longer evolutionary timescales. Hence, given the timeframe of our study, and the strong phylogenetic heritability in our data (see also [18]), it seems doubtful that the hypothesis can explain the evolvability-rate correlations observed for fly wings [19].

What remains surprising, then, is the conserved alignment between **D** (or **M**) and the observed divergence in absence of genetic constraints (Fig. 3C). An alternative explanation for our findings is that the observed patterns reflect the forces of stabilizing correlational selection exercising a similar influence on both developmental architectures and species divergence [16, 20, 22, 63]. To what extent genetic architecture and developmental systems can evolve by natural selection is, however, still a controversial question in need of further theoretical and empirical attention. For example, some recent models highlight that correlational selection can reshape **M** on relatively short time-scales in specific scenarios [11, 34, 35, 74, 75], while other models suggest that mutational and developmental biases evolve to align with the forces of correlational selection under fairly restrictive conditions [25, 26, 33, 76]. The main quandaries here are i) whether correlational selection on epistatic interactions can be strong and variable enough among traits to cause the relatively pronounced mutational and developmental biases often observed in quantitative traits, and ii) if such genetic architectures can be maintained for long enough to stably align with repeated macroevolutionary adaptations. Theory nevertheless suggests that conditions are particularly permissive when selection acts on multiple correlated characters (e.g., [77]) and fluctuates predictably between alternative fitness optima (e.g. [78]), which we argue is likely the case for patterns of selection on wing shape allometry, both within and between dipteran species (Fig. 6). Interestingly, allometries are often referred to as examples of constrained evolution based on the relatively stable nature of allometric exponents, and wing shape is no exception [79–81]. Yet, both theory and data also highlight that allometric relationships may be outcomes of common forces of correlational selection (e.g. for morphology [39], metabolism [82], growth [83], and reproductive investment [84]; but see [80] for an alternative interpretation regarding wing shape). Further work is needed to understand and describe correlational selection on allometric scaling in fly wings.

What is then the most plausible explanation for the observed correlations between measures of developmental bias, evolvability and macroevolutionary rates? Judging from the recent flurry of comparative studies [16, 18–20, 29], the answer seems to depend on the timescale and trait under consideration. For fly wing evolution on these large timescales, there has been little evidence for genetic constraints and a role for correlational selection simultaneously driving **D**, **G**, and **R** seems more plausible. However, explanations invoking genetic constraints or natural selection are not mutually exclusive and might simultaneously contribute to the alignment between developmental bias, evolvability, and evolution. Our study highlights the conundrum of explaining how these evolvability-rate relationships can persist over such deep macroevolutionary time (see also [20, 29, 39]). To understand the fundamental limits of adaptive morphological evolution, future studies must assess the theoretical plausibility of alternative explanations and identify the relevant timescales on which they apply. If developmental biases are indeed shaped by past forces of natural selection, then contemporary rates of adaptive evolution will depend on whether current selection pressures reflect those of the past, and when not, to what extent contemporary selection restructures the genotype-phenotype map and facilitates adaptation to new trait optima.

## Methods

### Quantifying wing shape and divergence across fly taxa

We focus on the evolution of wing shape within the Eremoneura, a clade within the Brachycera characterized by the presence of three larval instars. This clade is about 185 My old [42] and includes the dance and long-legged flies (Empidoidea) as well as the Cyclorrhapha (flies that pupate within the cuticle of the last larval instar (i.e., the puparium) [42]). We use the phylogenetic relationship among families proposed by Bayless et al. [47] as backbone for our comparative analysis.

To quantify the morphological variation within and between families, we took advantage of illustrations and pictures of fly wings from the taxonomic and systematic literature. An initial data set was sourced from the *Manuals of Nearctic Diptera* and the *Manuals of Afrotropical Diptera* [44, 85]. We focused on those families that are represented in the phylogenetic hypothesis generated by [47]. Additional pictures and illustrations were collected from a wide range of publications (see Supplementary Data Table 2).

Wing vein reduction has evolved numerous times across the phylogeny and can even be present as intraspecific (genetic) polymorphisms [86, 87]. Because homology is difficult to establish in these cases, we were unable to include these species in our analysis, in which we only included observations where the location of all 11 two-dimensional landmarks used in Rohner & Berger[16] could be assigned. In total, we collected wing shape data from 827 individuals belonging to 53 families. Using tpsDig2 [88], we manually quantified wing morphology as depicted on the illustrations and images. Additional morphometric data originally collected from images was added for 119 species of drosophilids from Houle et al. [18] and 36 sepsid species from [16]. The final data set contained 993 observations of 933 species in 530 genera and 68 families. Family affiliation of individual genera was checked using *Systema Dipterorum* [89].

The number of observations varied strongly across families (mean = 17.40, median = 12, minimum = 1, maximum = 130). This uneven sampling was caused by i) a varying number of species per family (e.g., Australimyzidae is a monogeneric family containing just 9 described species compared to Tachinidae with 9,626 species [90]), the loss of landmarks in several species (e.g., Sphaeroceridae [87]), and often incomplete illustrations or pictures showing only part of the wing (e.g., Muscidae). The landmark coordinates were aligned to the mean configuration of Houle et al. [18] using Procrustes analysis in MorphoJ [91].

To illustrate the main axes of morphological variation, we applied a Canonical Variate Analysis (CVA) in the R-package MASS [92]. This ordination technique finds the axes that maximize variation among species. For the CVA, we only considered those families with 5 or more individual observations.

### Estimating the phylogenetic variance-covariance matrix

The phylogeny includes the placement of individual families. We thus use the observation of different species within these families as repeated measures to approximate the phenotypic variation within the families. To calculate evolutionary rates per My, we calibrated the phylogeny by [47] using the R-package ape [93], modeling correlated substitution rate variation among branches. The approximate age of the Eremoneura (185 My [42]) and the divergence between Drosophilidae and Muscidae and Tephritidae (estimated to be 29-80 My and 48-86 My, respectively [94]) were used as calibration points. We computed the phylogenetic variance-covariance matrix **R** based on the inverse of the relationship matrix among families (S^-1^, [45]) using animal models in ASReml-R. To account for variation due to repeated observations within each family, we added species as an additional random effect (using ‘ide()’ structure). The samples within families thus serve as replicated measures. We only included families for which we had at least 5 species in our dataset.

### Estimating the dimensionality of covariance matrices

Although all our analysed matrices contain 18 dimensions (due to the loss of four dimensions for scaling, rotation, and positioning during Procrustes analysis), geometric morphometric covariance matrices are often rank deficient due to redundant covariance among landmark variables [95]. To assess how many dimensions of **R** had statistical support, we fitted reduced-rank factor analytic mixed models in ASReml-R. We began by fitting a covariance model with a single dimension and continually increased the number of dimensions until increasing the number of dimensions did not lead to a significant increase in model fil (based on Akaike’s Information Criterion (AIC)). We then extracted the reduced rank variance-covariance matrices from these best fitting models for further analysis using the R package ASExtra4 [96]. Error variances were estimated separately for each shape variable in all models.

### Comparison of variance-covariance matrices

The **D**, **G**, and **P** matrices for sepsids were taken from [16]. In brief, that study estimated **G** and **P** matrices based on a common garden experiment with 71 isofemale lines deriving from 7 populations of *S. punctum* and 42 lines and 9 populations of *S. fulgens*. The **D** matrix was calculated based on 87 male *S. punctum* and 96 male *S. fulgens*. **M**, estimated in its homozygous state in *Drosophila*, was extracted from [18]. Even though we focus on comparisons between **D** in *S. punctum* and our other variance-covariance matrices of interest, we compare the variances of these matrices along the eigenvectors of **P** estimated in *S. fulgens* to minimize bias in estimates of regression slopes [48]. Following [48, 49], we decomposed the **P** matrix estimated in *S. fulgens* into its eigenvectors and calculated the variance along for each of the respective variance-covariance matrices of interest as the diagonal entries of the matrix, where T denotes transposition and **X** refers to the matrix being compared (**D** and **G** for *S. punctum*; **M** for *D. melanogaster;* **R** for all Diptera measured). We then calculated Pearson’s correlation coefficients (r) and OLS slopes (b) between these logarithmized variances for a given pair of matrices. To avoid comparing matrices along null spaces with deficient variance, each matrix pair was compared along only the first *k* dimensions of **P,** with *k* equal to the rank of the matrix with lowest rank (10 dimensions in all cases).

To provide 95% confidence limits around correlations and slopes, we resampled the variance-covariance matrices from the factor analytical models with best support (based on AIC), using the REML-MVN approach [97]. This approach uses asymptotic resampling of REML estimates, taking advantage of the fact that the sampling distribution of variance-covariance matrices are well approximated by a multivariate normal distribution at large sample size. We performed the MVN resampling on the “G-scale” using the *mvtnorm* package for R [also see: 66, 67]. With this approach we resampled 10,000 matrices of each kind and subjected them to the common subspace analysis.

### Quantifying variation in genetic quality to test for deleterious pleiotropy associated with wing shape

To quantify fitness variance across the same isofemale lines of *S. punctum* as measured for wing shape, we reared all 71 lines (originating from the 7 European populations) in a common garden experiment including 9 temperature treatments ranging from 15-31°C. Note that these isofemale lines were created by pairing a single male and female and expanding population size to 100-200 flies immediately over a single generation and then maintaining lines at ca. N = 200 for 5-10 generations before the experiment. Thus, the studied among-line difference is likely to reflect more dominance variance than expected in a natural population due to inbreeding, but not to any extreme extent due to the rapid population expansion [98]. F0 containers with fly cultures were equipped with vials of previously frozen cow dung to attain freshly laid eggs. Each line was seeded with four vials per temperature treatment. For each vial, juvenile development rate was estimated as the inverse of the time (in days) between the date of a laid clutch and the subsequent emergence of F1 adults. Juvenile survival was calculated at the fraction of laid eggs that emerged as adults. Emerged F1 females were paired with a male from the same line and placed in a 50ml vial with access to sugar, water, and cow dung as egg laying substrate. Sugar, water, and dung were replaced every 5 days for the first 15 days. Early reproductive success was estimated as the total number of offspring produced within the first 15 days of adult female life, excluding females that died during this timeframe (likely due to accidental deaths). Females that did not lay any eggs during this period were not included in estimates of early reproductive success. Lifespan was estimated as the time from the start of the experiment until the focal female died. 245 females (17%) did not die during the period of observation and were recorded as censored data. After their death, females were measured for their tibia length as an estimate of body size [99]. In total, 1,445 females were measured across all lines and temperature treatments. These females produced a total of 173,556 offspring.

To test for heritable variation in early reproductive success, juvenile survival, developmental rate, and body size, we used mixed effects models using restricted maximum likelihood as implemented in *ASReml-R* [46]. Temperature treatment, population, as well as their interaction were fitted as fixed effect. Line was added as random effect. Note that we did not estimate line by treatment interactions (i.e. G-by-E) as our aim was to capture overall differences in genetic quality among lines. Repeating the analysis excluding the highest (31□) and lowest (15□) (i.e. most stressful) temperature treatments led to similar results. Residual variances were allowed to vary across treatments. The significance of the random effect of line was tested using Likelihood Ratio Tests (LRTs). Best linear unbiased predictors (BLUPs) were extracted and used for further analysis. For the analysis of adult lifespan, we fitted a censored mixed effects cox model using the *coxme* package [100] using treatment and population as fixed effects and line as random effect. LRTs were used to test whether line effects were significant. BLUPs were extracted by taking the inverse of the hazard ratio for each line as our estimates for adult longevity.

We applied principal component analysis on the correlation matrix based on BLUPs to inspect the loadings on PC1. To test whether the proportion of variance explained by PC1 is larger than expected by chance we used two different randomization procedures. First, we simulated 10,000 random and unstructured covariance matrices based on the same sample size as in our real data (using the R function *rnorm()*) to generate a null distribution for the proportion of variance explained by the first PC. Secondly, we generated an alternative null distribution by randomizing breeding values among families 10,000 times and calculated the relative eigenvalue of the first eigenvector that was compared with the observed eigenvalue for PC1.

To test for genetic correlations between wing shape and fitness, we used two-block partial least squares regression as implemented in *geomorph* [101]. This is an ordination technique that finds the latent variables within two sets of variables with maximal covariance between the two sets of variates. We used wing shape in the first block. To account for shape allometry and local adaptation, we used the residuals of a multivariate regression of shape on centroid size and population. The second block either consisted of scores on PC1 based on the five fitness correlates, or breeding values for the five variables separately. Significance was assessed using permutation tests (10’000 random permutations).

### Testing for constraints: estimating the relationship between the rate and direction of evolution with respect to **D**

To test whether evolutionary change that aligns with **D** is faster than evolutionary change in other directions (as expected if **D** constrains macroevolution), we reconstructed ancestral wing shape at each node and extracted the vectors of shape change observed on each edge of the phylogeny (using the *gm.prcomp()* function in *geomorph*). We then quantified the developmental variance captured by shape evolution as:

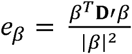

where *β* is the evolutionary shape change vector of interest, **D’** is the **D** matrix scaled by its trace, and *T* denotes transposition. If evolution occurs primarily along the main axes of **D**, we expect *e*_β_ to be large and close to the variance explained by the first eigenvector of **D** (d_max_). In contrast, if shape evolution is independent of **D**, this variance is expected to be small and closer to the variance explained by the tenth eigenvector of **D** (d_10_). To test for constraints, we computed the correlation between *e*_β_ and the magnitude of the shape change along all edges in the phylogeny (measured as shape change in Procrustes distance per My), expecting a positive correlation if fast rates of evolution are constrained to occur along dimensions with high developmental variability. These analyses were repeated using **M** as base of comparison.

To compare the observed correlation between the rate of evolution and its direction with respect to **D**, to the expected correlation under Brownian motion, we simulated wing shape evolution using the *mvSIM* function implemented in the *mvMORPH* package [102]. To simulate Brownian motion, where the evolutionary changes are more likely to occur along the dimensions of high evolvability, we sampled evolutionary changes from a distribution *N*(μ, Σ), where μ is a 22-dimensional vector of means equal to zero and Σ is a covariance matrix set equal to **D** scaled to the same size as **R**. In these simulations, evolutionary changes are thus governed by Brownian motion but constrained by the orientation of **D**. For each simulation, we then computed *e*_β_ as described above and compared the resulting distribution of *e*_β_ and its correction with **D** to the observed values. We again repeated this analysis using the **M** matrix.

### Testing for allometry

To assess whether allometric scaling is associated with **D**/**M**, we calculated centroid size for those specimens where scale bars were available (n = 127). We then used a multivariate regression of wing shape on log centroid size to estimate the evolutionary allometric shape change vector (using procD.lm in geomorph). A phylogenetic regression of family mean shape on mean centroid size was also significant and resulted in a very similar vector (vector correlation: r = 0.92, P <.001). The alignment between this vector and **D** and **R** was calculated using the method described above. Allometric shape changes were visualized using shape scores following [103].

