## Supplementary Data for "Macroevolution of fly wings proceeds along developmental lines of least resistance"

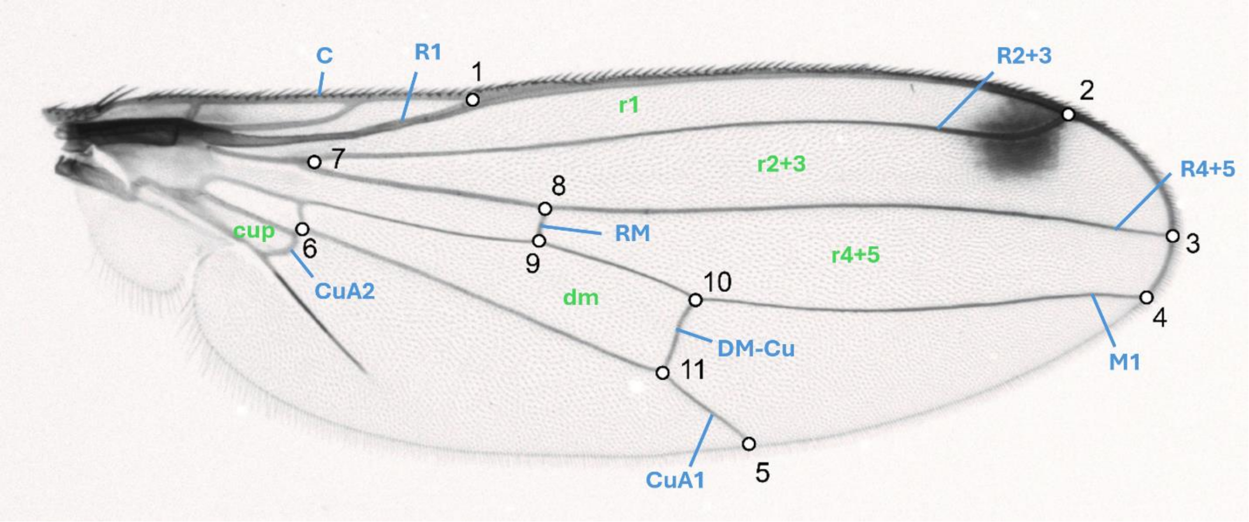


**Supplementary Data Figure 1:** Location of two-dimensional landmarks used for this study illustrated on a picture of a sepsid wing. Wing veins are indicated in blue. Cells are indicated in green. Nomenclature follows [1] (also see [2]).

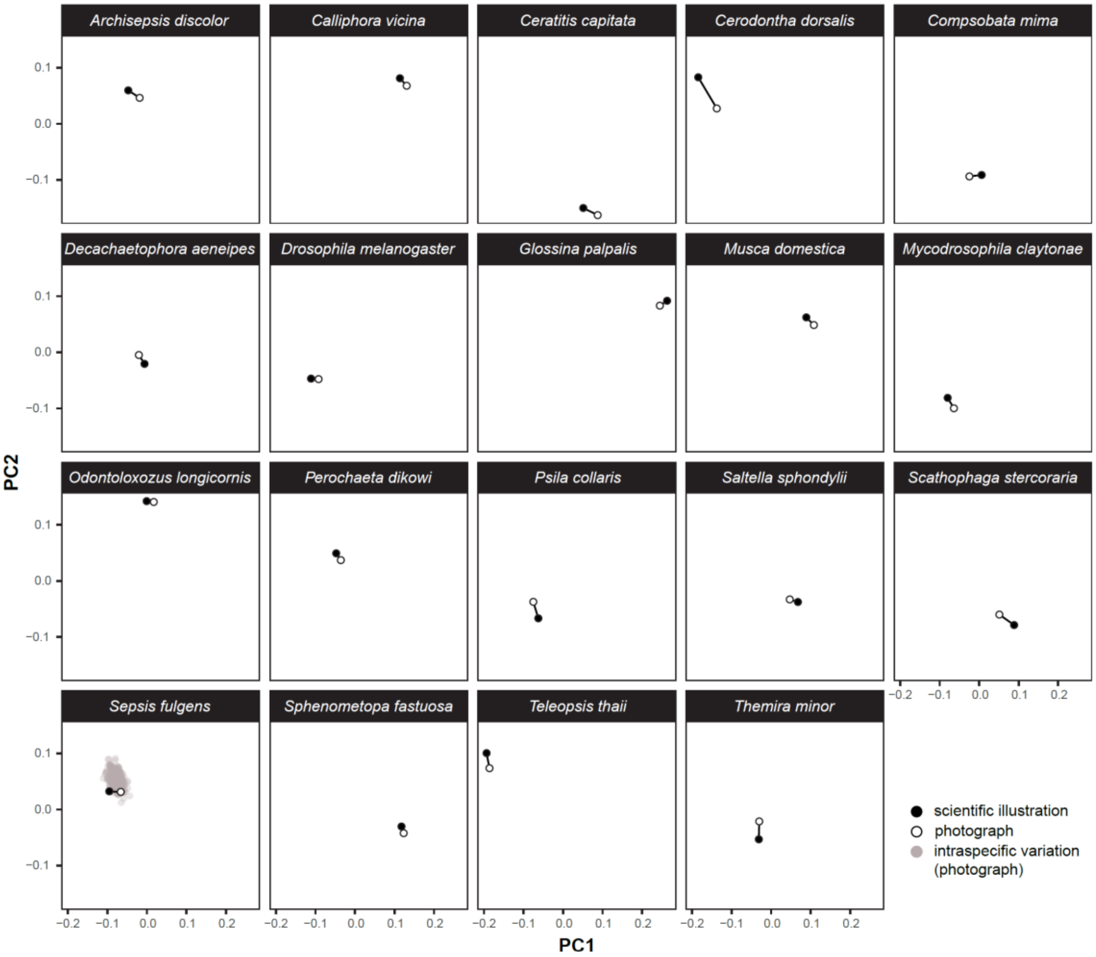


**Supplementary Data Fig. 2:** Morphospace (principal components) showing the similarity between wing shape measured on photographs or scientific illustrations from the systematic and taxonomic literature. The intraspecific variation observed in *Sepsis fulgens* (data from [3]) was projected into the morphospace to provide a reference point for the amount of morphological disparity that can be expected in natural populations.

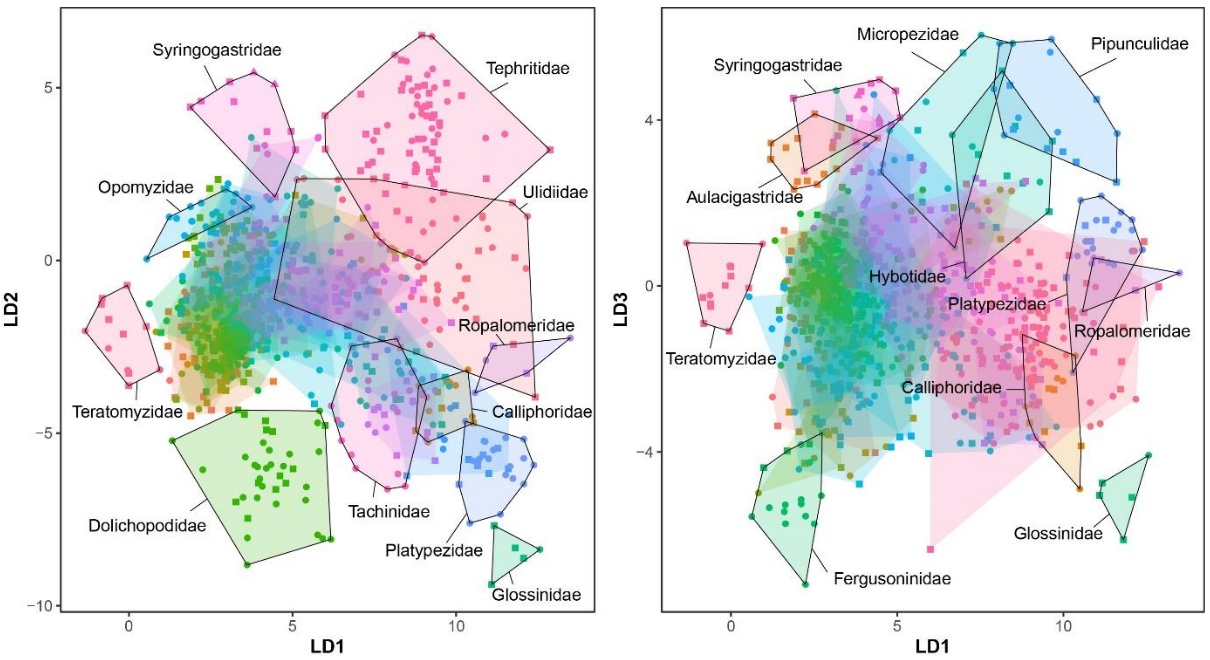


**Supplementary Data Fig. 3:** Evolutionary morphospace defined by the first three canonical variates. Individuals are grouped by family (hulls). An arbitrary set of families is highlighted. The source of the shape data is indicated with shape (scientific illustration and pictures). Families with less than five observations were excluded from the analysis, leading to a total number of 43 families (n = 884). The overall leave-one-out cross-validation success of the CVA was 82.3%.

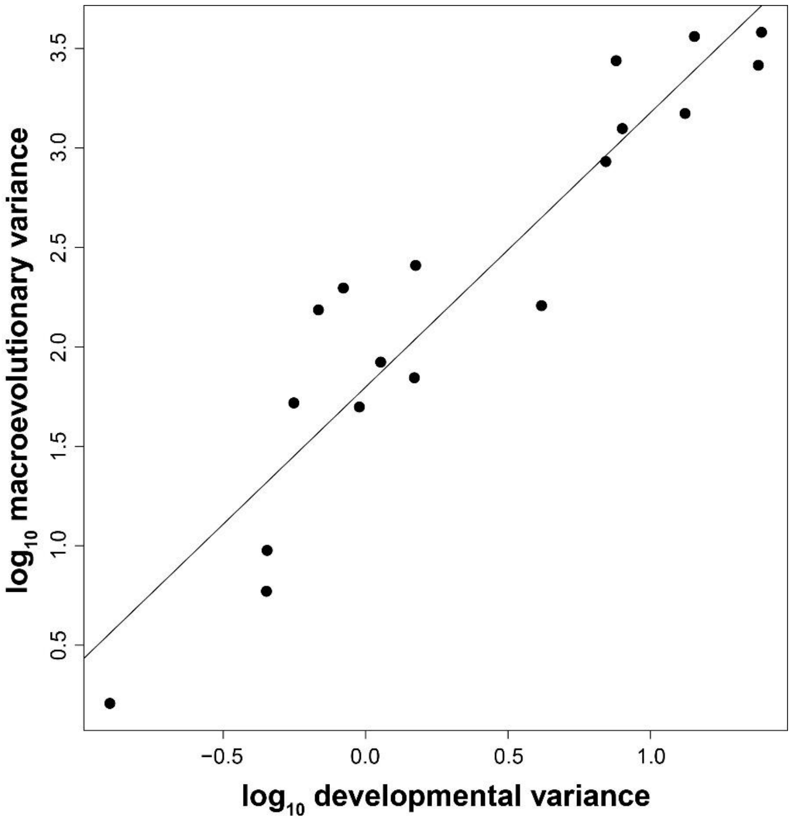


**Supplementary Data Fig. 4:** Results of a common subspace analysis where the amount of developmental variance predicts the macroevolutionary variance along all possible 18 eigenvectors of the **P** matrix estimated in *S. fulgens* (slope = 1.38, r = 0.94).

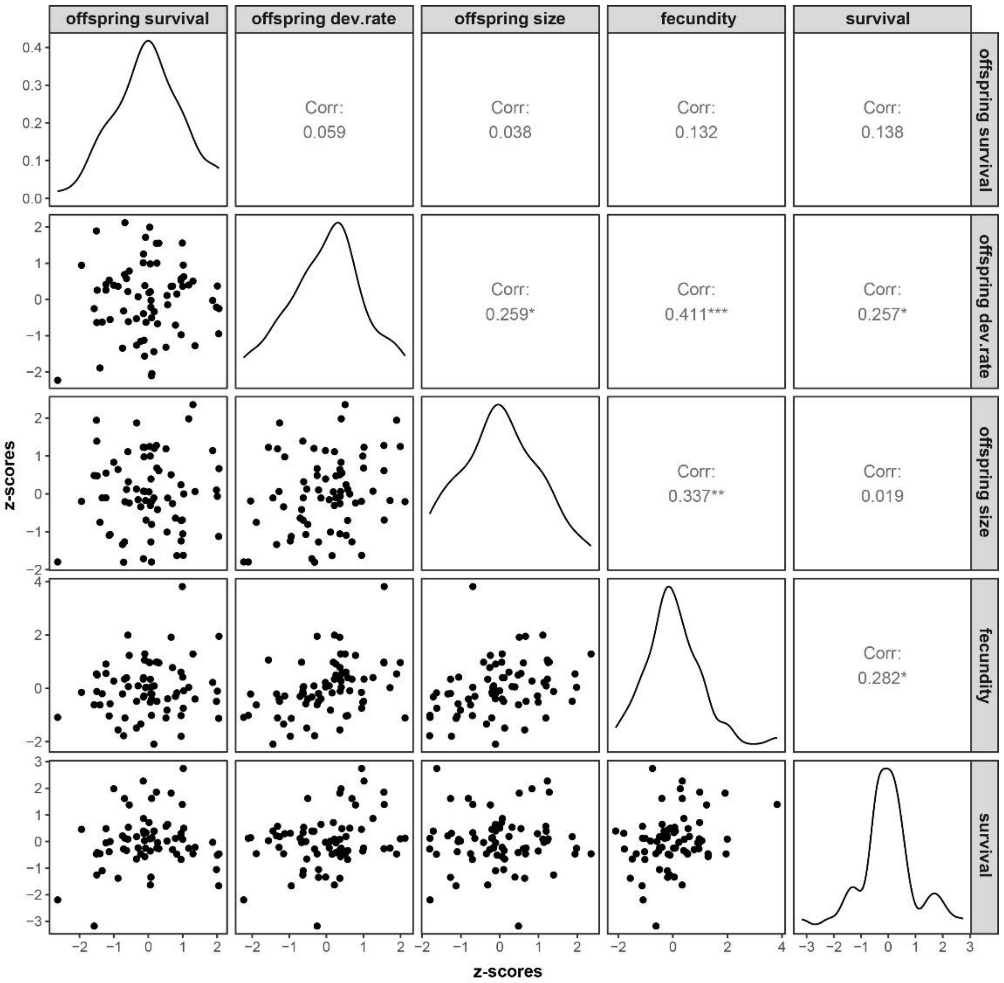


**Supplementary Data Fig. 5:** Genetic correlations between fitness components. Data points indicate best linear unbiased predictions (BLUPs). BLUPs for offspring survival, developmental rate, size and adult fecundity were extracted from mixed effect models using restricted maximum likelihood. BLUPs for adult lifespan were extracted from a censored mixed effects cox model. All data have been transformed to z-scores.

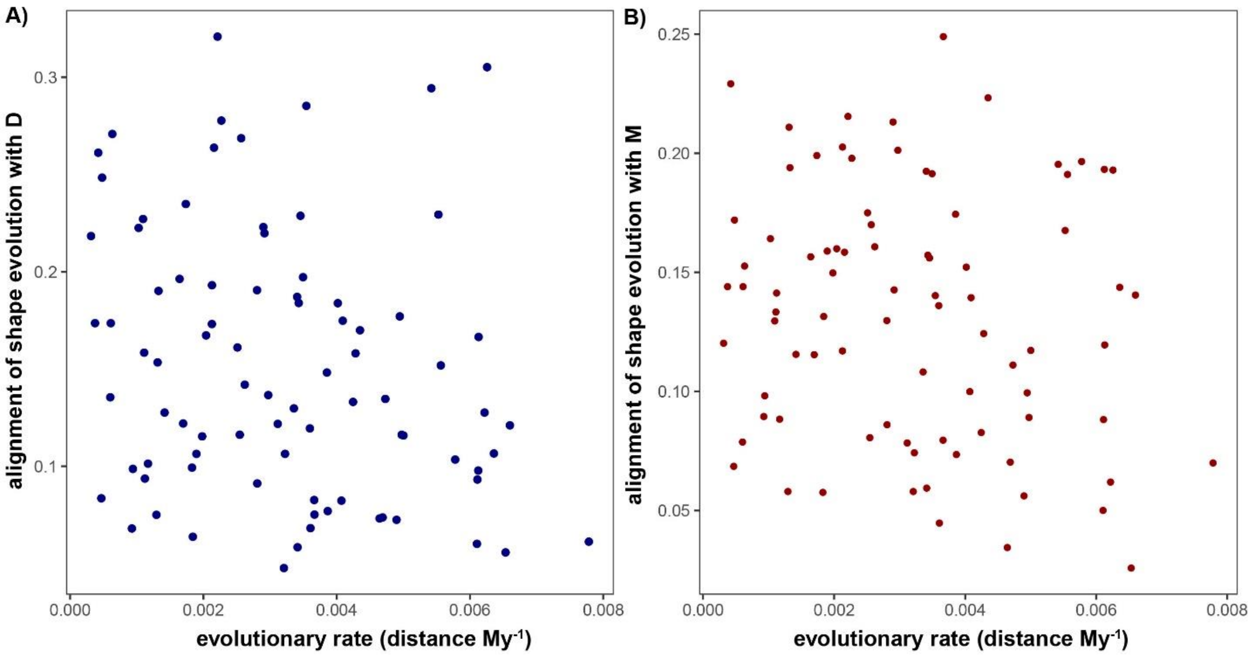


**Supplementary Data Fig. 6:** The relationship between evolutionary rate (in Procrustes distance per My) along individual branches of the phylogeny and the amount of variance in either matrix captured by the vector of evolutionary shape change. In contrast to the expectation based on the constraint hypothesis, the speed of evolutionary change does not increase when its direction aligns with the main axes of either matrix.

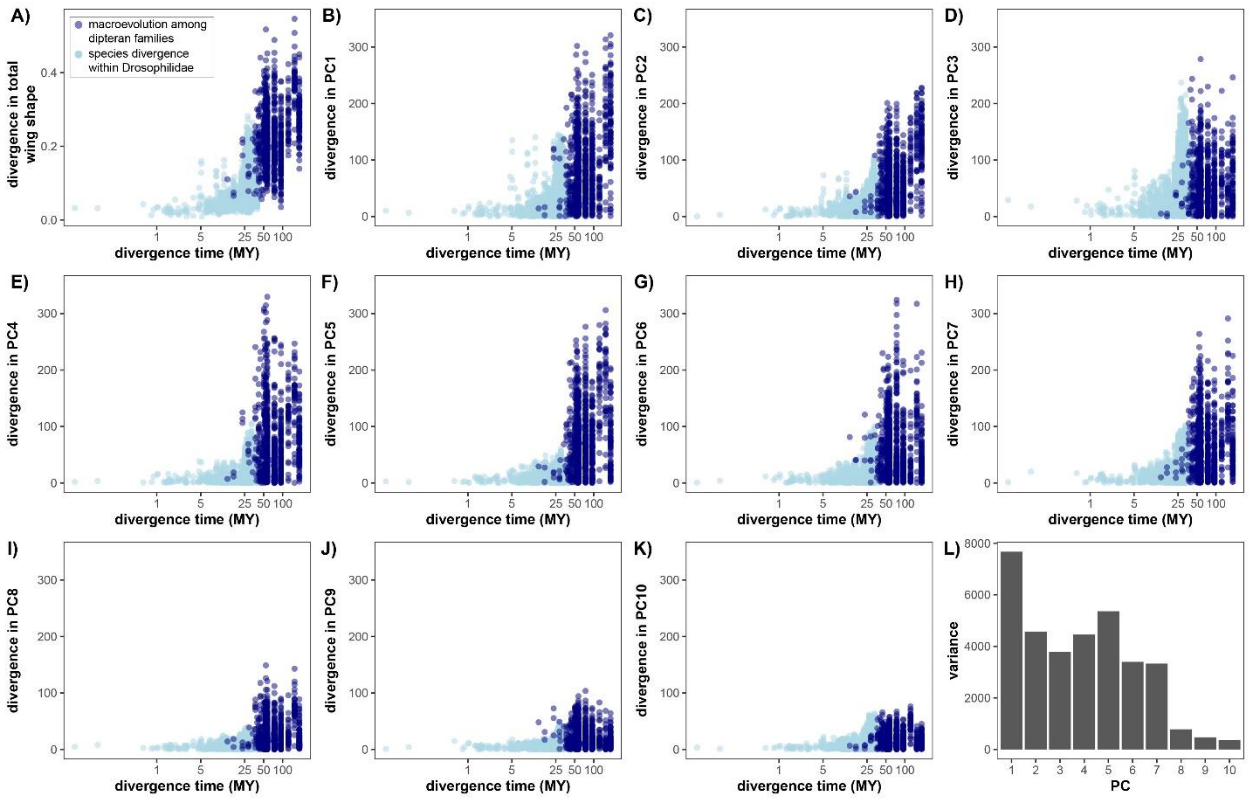


**Supplementary Data Fig. 7:** Accumulation of divergence in overall wing shape (quantified by Procrustes distance, A), as well as the first 10 principal components with evolutionary time (B-K). For comparison with the common subspace analysis, the wing shape data were first multiplied by 1,000 and then projected onto the eigenvectors of the phenotypic variance covariance matrix (**P**) estimated in *Sepsis fulgens (*as in the main analyses presented in the main text*)*. Panel L) shows the total amount of variance in our dataset along the first ten eigenvectors of **P**. Data on species divergence among drosophilids (spanning up to 40 My) are from [4].

**Supplementary Data Table 1:** Landmarks were defined based on the position of the longitudinal wing veins (Costa (C), Radius (R1-5), Media (M1-3), Cubitus (CuA1)) and the two cross-veins R-M and DM-Cu, as well as the wing cells that are delineated by these veins (see Supplementary Data Figure 1). Nomenclature follows [1] (also see [2]).

| Landmark | morphological description |
|---|---|
| LM1 | intersections between R1 and C |
| LM2 | intersections between R2+3 and C |
| LM3 | intersections between R4+5 and the wing margin |
| LM4 | intersections between M1 and the wing margin |
| LM5 | distal end of CuA1 |
| LM6 | connection between cell cup and CuA1 |
| LM7 | proximal tip of cell r2+3 (branching of R2+3 and R4+5) |
| LM8 | intersection between R-M and R4+5 |
| LM9 | intersection between R-M and M1 |
| LM10 | intersection between DM-Cu and M1 |
| LM11 | intersection between DM-Cu and CuA1 |

**Supplementary Data Table 2:** Sources of illustrations and pictures of fly wings, including total sample sizes (n).

| taxon | source | n |
|---|---|---|
| Acartophthalmidae | [5-7] | 4 |
| Agromyzidae | [5, 8-14] | 16 |
| Anthomyzidae | [12, 15-18] | 25 |
| Asteiidae | [11, 12, 19] | 4 |
| Atelestidae | [2] | 1 |
| Aulacigastridae | [20] | 15 |
| Australimyzidae | [21] | 1 |
| Calliphoridae | [12, 22-29] | 14 |
| Canacidae | [11, 12] | 8 |
| Carnidae | [11, 12] | 3 |
| Chamaemyiidae | [5, 11, 12, 30, 31] | 12 |
| Chloropidae | [5, 32] | 3 |
| Chyromyidae | [12, 33, 34] | 11 |
| Clusiidae | [5, 11, 12] | 13 |
| Coelopidae | [5, 11, 12, 34-36] | 10 |
| Cryptochetidae | [5, 30, 34, 37-39] | 10 |
| Curtonotidae | [5, 11, 40] | 17 |
| Diastatidae | [11, 12, 41] | 17 |
| Diopsidae | [12, 42-45] | 31 |
| Dolichopodidae | [2, 5, 11, 46] | 39 |
| Drosophilidae | [4, 5, 12, 30] | 130 |
| Empididae | [2, 46, 47] | 3 |
| Ephydridae | [5, 12, 48] | 12 |
| Fergusoninidae | [31, 44, 49-54] | 16 |
| Glossinidae | [55-59] | 5 |
| Heleomyzidae | [11, 12, 60-64] | 30 |
| Hybotidae | [2, 46] | 12 |
| Lauxaniidae | [5, 11, 12] | 31 |
| Micropezidae | [12, 44, 65, 66] | 12 |
| Muscidae | [12, 67-71] | 25 |
| Neriidae | [5, 11, 12, 44, 65, 66, 72-74] | 11 |
| Odiniidae | [5, 12, 75-81] | 21 |
| Opomyzidae | [11, 12, 82] | 28 |
| Paraleucopidae | [12, 83] | 2 |
| Periscelididae | [5, 11, 12, 34, 84-89] | 16 |
| Piophilidae | [5, 11, 12, 90, 91] | 14 |
| Pipunculidae | [5, 11, 12, 30, 55] | 15 |
| Platypezidae | [5, 11, 12] | 22 |
| Psilidae | [5, 11, 12, 44, 92] | 11 |

| | | |
|---|---|---|
| Ropalomeridae | [5, 93, 94] | 5 |
| Sarcophagidae | [12, 30, 95-101] | 20 |
| Scathophagidae | [12, 102-104] | 11 |
| Sciomyzidae | [11, 12, 105, 106] | 30 |
| Sepsidae | [5, 12, 107-109] | 38 |
| Sphaeroceridae | [11, 12, 110] | 8 |
| Strongylophthalmyiidae | [44, 111] | 6 |
| Syringogastridae | [44, 112, 113] | 11 |
| Tachinidae | [12, 55, 114] | 17 |
| Tephritidae | [11, 12, 115] | 72 |
| Teratomyzidae | [116, 117] | 13 |
| Ulidiidae | [5, 11, 12, 115, 118-120] | 47 |

**Supplementary Data Table 3:** Leave-one-out cross validation success of the Canonical Variate Analysis (CVA). Only families with 5 or more observations were considered (n = 884).

| | sample size | cross validation success |
|---|---|---|
| Agromyzidae | 16 | 0.75 |
| Anthomyzidae | 25 | 0.96 |
| Aulacigastridae | 14 | 1.00 |
| Calliphoridae | 13 | 0.77 |
| Canacidae | 8 | 0.25 |
| Chamaemyiidae | 12 | 0.75 |
| Chyromyidae | 11 | 1.00 |
| Clusiidae | 12 | 0.75 |
| Coelopidae | 10 | 0.80 |
| Cryptochetidae | 10 | 1.00 |
| Curtonotidae | 17 | 0.82 |
| Diastatidae | 16 | 0.81 |
| Diopsidae | 31 | 0.90 |
| Dolichopodidae | 39 | 0.87 |
| Drosophilidae | 130 | 0.92 |
| Ephydridae | 11 | 0.55 |
| Fergusoninidae | 16 | 1.00 |
| Glossinidae | 5 | 1.00 |
| Heleomyzidae | 6 | 0.17 |
| Hybotidae | 12 | 1.00 |
| Lauxaniidae | 30 | 0.60 |
| Micropezidae | 11 | 0.91 |
| Muscidae | 22 | 0.59 |
| Neriidae | 11 | 0.64 |
| Odiniidae | 21 | 0.86 |
| Opomyzidae | 27 | 1.00 |
| Periscelididae | 16 | 0.31 |
| Piophilidae | 13 | 0.38 |
| Pipunculidae | 15 | 1.00 |
| Platypezidae | 22 | 1.00 |
| Psilidae | 10 | 1.00 |
| Ropalomeridae | 5 | 1.00 |
| Sarcophagidae | 19 | 0.68 |
| Scathophagidae | 10 | 0.70 |
| Sciomyzidae | 29 | 0.72 |
| Sepsidae | 38 | 0.97 |
| Sphaeroceridae | 8 | 0.88 |
| Strongylophthalmyiidae | 6 | 1.00 |
| Syringogastridae | 11 | 1.00 |
| Tachinidae | 16 | 0.63 |
| Tephritidae | 71 | 0.97 |
| Teratomyzidae | 13 | 1.00 |
| Ulidiidae | 46 | 0.74 |

**Supplementary Data Table 4**: A modified version of Krzanowski's common subspace analysis [4, 121, 122] was used to compare different covariance matrices along an arbitrary set of orthogonal phenotypic dimensions (or eigenvectors). Because the choice of reference matrix can bias estimates of slopes and correlation coefficients, all comparisons were repeated using different sets of eigenvectors. *k* refers to the number of dimensions considered for comparisons.

**A) eigenvectors used for comparison: phenotypic variation (P) in *S.punctum***

| matrix compared to **R** | *k* | slope [95% CI] | r [95% CI] |
|---|---|---|---|
| **D** (*S. punctum*) | 10 | 0.49 [0.4, 0.57] | 0.85 [0.76, 0.91] |
| **D** (*S. fulgens*) | 9 | 0.7 [0.56, 0.81] | 0.93 [0.84, 0.97] |
| **G** (*S. punctum*) | 10 | 0.96 [0.83, 1.05] | 0.87 [0.8, 0.91] |
| **G** (*S. fulgens*) | 10 | 0.79 [0.67, 0.9] | 0.92 [0.84, 0.95] |
| **P** (*S. punctum*) | 10 | 0.85 [0.74, 0.94] | 0.91 [0.85, 0.94] |
| **P** (*S. fulgens*) | 10 | 0.69 [0.59, 0.78] | 0.89 [0.82, 0.93] |
| **M** (*D. melanogaster*) | 10 | 0.57 [0.46, 0.67] | 0.81 [0.69, 0.88] |
| **G** (*D. melanogaster*) | 10 | 0.56 [0.48, 0.61] | 0.88 [0.81, 0.92] |
| **R** (*D. melanogaster*) | 10 | 0.75 [0.64, 0.82] | 0.83 [0.76, 0.88] |

**B) eigenvectors used for comparison: phenotypic variation (P) in *S.fulgens***

| matrix compared to **R** | *k* | slope [95% CI] | r [95% CI] |
|---|---|---|---|
| **D** (*S. punctum*) | 10 | 0.61 [0.5, 0.71] | 0.87 [0.78, 0.92] |
| **D** (*S. fulgens*) | 9 | 0.43 [0.32, 0.53] | 0.82 [0.67, 0.9] |
| **G** (*S. punctum*) | 10 | 0.9 [0.75, 1.01] | 0.85 [0.76, 0.9] |
| **G** (*S. fulgens*) | 10 | 0.95 [0.79, 1.08] | 0.93 [0.86, 0.97] |
| **P** (*S. punctum*) | 10 | 0.79 [0.66, 0.89] | 0.87 [0.78, 0.92] |
| **P** (*S. fulgens*) | 10 | 0.81 [0.68, 0.9] | 0.88 [0.8, 0.93] |
| **M** (*D. melanogaster*) | 10 | 0.66 [0.53, 0.77] | 0.89 [0.78, 0.94] |
| **G** (*D. melanogaster*) | 10 | 0.53 [0.43, 0.61] | 0.78 [0.68, 0.85] |
| **R** (*D. melanogaster*) | 10 | 0.36 [0.26, 0.47] | 0.48 [0.35, 0.6] |

**C) eigenvectors used for comparison: developmental variation (D) in *S.fulgens***

| matrix compared to **R** | *k* | slope [95% CI] | r [95% CI] |
|---|---|---|---|
| **D** (*S. punctum*) | 10 | 0.58 [0.49, 0.66] | 0.82 [0.74, 0.88] |
| **D** (*S. fulgens*) | 9 | 0.44 [0.31, 0.56] | 0.67 [0.52, 0.79] |
| **G** (*S. punctum*) | 10 | 0.89 [0.78, 0.98] | 0.89 [0.84, 0.93] |
| **G** (*S. fulgens*) | 10 | 0.99 [0.87, 1.09] | 0.94 [0.89, 0.96] |
| **P** (*S. punctum*) | 10 | 0.88 [0.78, 0.96] | 0.96 [0.92, 0.98] |
| **P** (*S. fulgens*) | 10 | 0.78 [0.69, 0.87] | 0.92 [0.86, 0.95] |
| **M** (*D. melanogaster*) | 10 | 0.47 [0.39, 0.54] | 0.78 [0.67, 0.86] |
| **G** (*D. melanogaster*) | 10 | 0.57 [0.5, 0.62] | 0.87 [0.81, 0.91] |
| **R** (*D. melanogaster*) | 10 | 0.62 [0.54, 0.68] | 0.87 [0.81, 0.91] |

**Supplementary Data Table 5:** Observed and expected amount of macroevolutionary variance along 10 phenotypic dimensions under a pure drift scenario. The expected divergence under drift was calculated as two times the mutational variance along each dimension times 1.85 $\times 10^{8}$ (this is equivalent to assuming one fly generation per year for 180MY). For consistency with Figs. 3 and S6, the dimensions 1-10 are the first 10 eigenvectors of the phenotypic variance covariance matrix (**P**) estimated in *S. fulgens*. In all instances, the observed macroevolutionary variance is more than 4,000 times smaller than expected under a pure drift scenario. Shape variables were multiplied by 1,000 (as in [4])

| dimension | observed macro-evolutionary variance | contemporary mutational variance | expected divergence under drift | ratio of expected to observed variance |
|---|---|---|---|---|
| **1** | $4.6 \times 10^{3}$ | $1.5 \times 10^{-1}$ | $5.5 \times 10^{7}$ | $1.2 \times 10^{4}$ |
| **2** | $2.7 \times 10^{3}$ | $1.1 \times 10^{-1}$ | $4.2 \times 10^{7}$ | $1.6 \times 10^{4}$ |
| **3** | $9.3 \times 10^{2}$ | $9.2 \times 10^{-2}$ | $3.4 \times 10^{7}$ | $3.7 \times 10^{4}$ |
| **4** | $2.6 \times 10^{3}$ | $5.2 \times 10^{-2}$ | $1.9 \times 10^{7}$ | $7.4 \times 10^{3}$ |
| **5** | $3.5 \times 10^{3}$ | $3.9 \times 10^{-2}$ | $1.5 \times 10^{7}$ | $4.2 \times 10^{3}$ |
| **6** | $1.8 \times 10^{3}$ | $5.5 \times 10^{-2}$ | $2.0 \times 10^{7}$ | $1.2 \times 10^{4}$ |
| **7** | $1.4 \times 10^{3}$ | $6.1 \times 10^{-2}$ | $2.3 \times 10^{7}$ | $1.6 \times 10^{4}$ |
| **8** | $2.8 \times 10^{2}$ | $1.7 \times 10^{-2}$ | $6.1 \times 10^{6}$ | $2.2 \times 10^{4}$ |
| **9** | $1.5 \times 10^{2}$ | $1.0 \times 10^{-2}$ | $3.7 \times 10^{6}$ | $2.4 \times 10^{4}$ |
| **10** | $8.5 \times 10^{1}$ | $7.2 \times 10^{-3}$ | $2.7 \times 10^{6}$ | $3.2 \times 10^{4}$ |
